# Epithelial-mesenchymal boundary guides cell shapes and axis elongation in embryonic explants

**DOI:** 10.1101/2024.08.20.608779

**Authors:** Katia Barrett, Shalabh Anand, Virginie Thome, Pierre-François Lenne, Matthias Merkel

## Abstract

The spatial arrangement of tissues during development establishes regions of tissue-tissue interactions. While such interactions are prominent during gastrulation and organogenesis, how they guide morphogenesis remains unclear. Here, using Xenopus animal cap explants, we show that the coupling between epithelial and mesenchymal tissues is instrumental in robust axis elongation. We find that mesenchymal and epithelial tissues drive elongation hierarchically - the epithelium can elongate independently, while the mesenchyme requires the initial presence of the epithelium. The epithelial-mesenchymal boundary defines the direction of cell shape alignment and the axis of tissue elongation. Prior to explant elongation, epithelial cells align and elongate, a process that propagates into the underlying mesenchyme. These findings reveal how tissue boundaries can organize coherent axis elongation without external patterning cues, providing insights into the self-organization principles that guide embryonic morphogenesis.

## Introduction

Embryonic morphogenesis is a complex process characterized by coordinated movements of tissues, driven by an interplay of mechanical and biochemical activities. Within this process, morphogenetic patterning leads to regional cell differentiation, which imbues cells with specific behaviors, giving rise to different tissues, which is crucial for shaping the developing organism. During morphogenesis, neighboring tissues interact and impose organizational constraints on one another. This coupling between neighboring tissues becomes particularly prominent after gastrulation, which establishes the three primary germ layers. During organ morphogenesis, differential growth between adjacent tissues can generate folded patterns and complex shapes, as seen in the intestine (Savin et al., 2011), lung (Varner & Nelson, 2015), and brain ((Tallinen et al., 2016)). There is also growing evidence that inter-tissue mechanical coupling is crucial for convergent extension (CE) mechanisms (Xiong et al., 2020), (Collinet 2015). Through this transformation, tissues extend preferentially along a specific spatial direction, thereby contributing to the overall elongation of the embryo or of individual organs. The textbook model of CE is largely influenced by the seminal work of R. Keller on Xenopus, which posits that the forces driving CE are generated by the deep mesenchymal tissues that polarise mediolaterally and intercalate beneath the presumptive mesodermal and neural tissues (Keller et al., 2008). Notably, the initial stages of this process are contingent upon the mesodermal mesenchymal cell attachment to the overlying endodermal epithelium, without which tissue extension fails (Wilson et al., 1989). The capacity for tissue extension is recovered when deep mesodermal cells remain coupled to the overlying endodermal epithelium, even outside the environment of the embryo. This implies that epithelial-mesenchymal tissue interaction is important for axial extension but how the elongation axis emerges, and to which extent the coupling between epithelial and mesenchyme tissues regulate this process remain unknown.

To tackle the questions, we focus here on Xenopus animal cap explants as a minimal model comprising two coupled tissues that can self-organize. Extracted from the pre-gastrula embryo, they are formed by pluripotent pre-ectodermal cells that can be pushed toward any germ layer. They are arranged as a bilayer comprising a single outer layer of epithelial cells and two inner layers of mesenchymal cells (Fig. 1A, epithelial cells in blue, mesenchymal cells in green). Upon removal from the embryo, the mesenchymal cells undergo a rounding-up process, while the epithelial layer crawls over the mesenchymal cells mirroring the natural process of epiboly. Addition of Activin A at a concentration of 10ng/ml to the culture medium triggers explant elongation (Fig. 1A-C, Movie 1) (Sawamura and Uchiyama, n.d.). This induced elongation replicates the behavior observed during axis extension in intact embryos, albeit with different geometric constraints.

**Figure 1.**
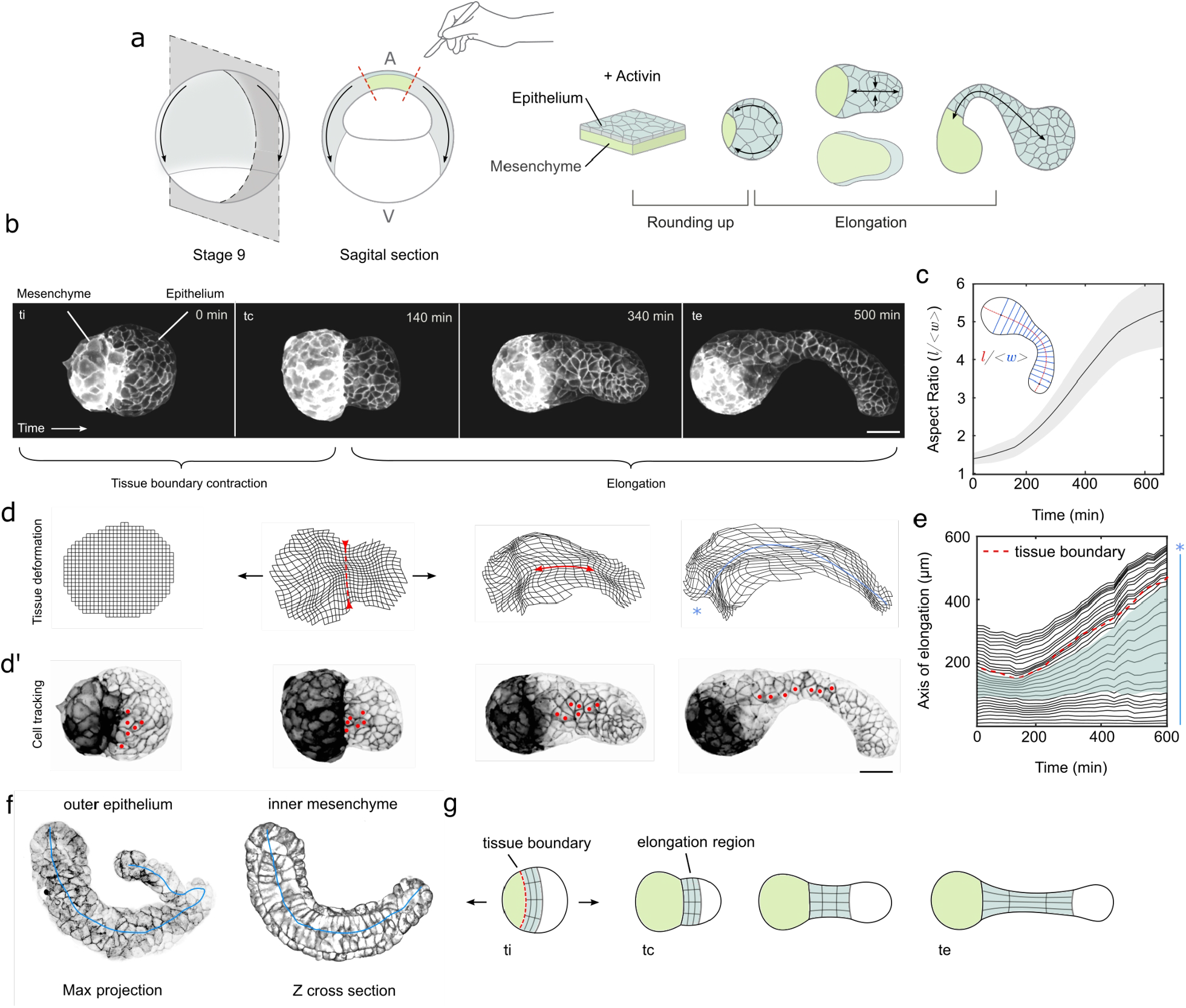
Xenopus animal cap explants display localized elongation. A) Schematic of the experimental protocol to induce elongation in animal cap explants. B) Snapshots of explant elongation with defining morphological timepoints. ti - initial timepoint, tc - tissue contraction, te - elongation. Scale bar = 100µm C) The aspect ratio is defined as the longest axis l divided by the averaged widths <w> measured at right angles to the elongation axis. Scale bar = 100 µm. N=8. D) Tissue deformation at different timepoints. D’) tracking a group of cells at the boundary of the explant. E) Kymograph of the elongation axis tissue deformation over time. Green shaded region highlights the region that is elongating, blue asterisk and line relate to the axis shown in D. F) Example of internal mesenchymal and outer epithelium in an elongated explant. G) Schematic of localised elongation reflecting tissue deformation kymograph.

## Results

### Activin-stimulated Xenopus animal cap explants display strong elongation

We first replicated classical experiments that demonstrate that animal cap explants are able to elongate on their own in the presence of Activin (Fig. 1A) (Symes and Smith, 1987), while they do not in its absence (Supplemental Fig. 1, Movie S2). We find that after 40 minutes, the initially roughly flat explants round up to an approximately spherical shape (Fig. 1B). After further 4 hours, the explant elongates over a time course of 10 hours along an axis perpendicular to the mesenchyme (Movie S1). We quantified the time-dependent aspect ratio as the length of the (curved) long axis *l* divided by the averaged width <*w>* (Methods), and found that our explants reached an aspect ratio of approximately 5 (Fig. 1C). We found that prior to elongation the epithelial region contracted but especially circumferentially around the epithelial-mesenchymal boundary (Fig. 1B, labels).

We noticed that CE does not occur homogeneously across the whole aggregate (Fig 1D, Movie S1). Instead, the epithelial cells closer to the mesenchyme-epithelial boundary rearrange strongly (Fig. 1D’ [marked rearranging cells], Movie S1), while at both mesenchyme and epithelial ends, we observe virtually no cell rearrangements. To quantify this, we applied particle image velocimetry to time-lapse images of the elongating aggregates (Methods, Fig. 1D [snapshots distorted square grid], Movie S1). We found that elongation exclusively localized to an initially narrow epithelial region close to the mesenchyme-epithelial boundary (Fig. 1E [plot]). Using fixed explants, we verified that this elongating region is made of the epithelium covering mesenchyme (Fig. 1F). Our observations show that tissue elongation results mainly from the CE of a narrow band of tissue near the edge of the epithelium, where it covers the mesenchyme (Fig. 1G).

### Only explants with an activin-stimulated epithelium are able to elongate

To dissect the respective roles of epithelium (epi) and mesenchyme (mes) in driving the elongation process, we leveraged the ease of the system, where it is possible to isolate tissues using micro-dissection and test how they behave separately when primed or not with Activin (Fig. 2A). Consistent with previous findings, mesenchymal cells alone, even when stimulated with Activin, failed to elongate: the mesenchymal tissue explants remained spherical (Fig 2B, B’ Movie S3). As did isolated Epithelium and Mesenchymal tissues without Activin (Movie S3, Supplemental Fig. S2). Epithelial tissue explants showed a much greater capacity to elongate in isolation but with more variability than the full explants (compare Fig. 2C, C’ to Fig. 1B,C, Movie S3), some elongating extensively, some with multiple poles and others just showing transient deviations from a spherical shape (Fig 2C, Movie S4 to be made). The fact that the epithelium elongates independently, demonstrates that the epithelium can create active oriented forces driving tissue extension. Yet, the observed variability also suggests that the contact of both epithelium and mesenchyme is required for robust elongation.

**Figure 2.**
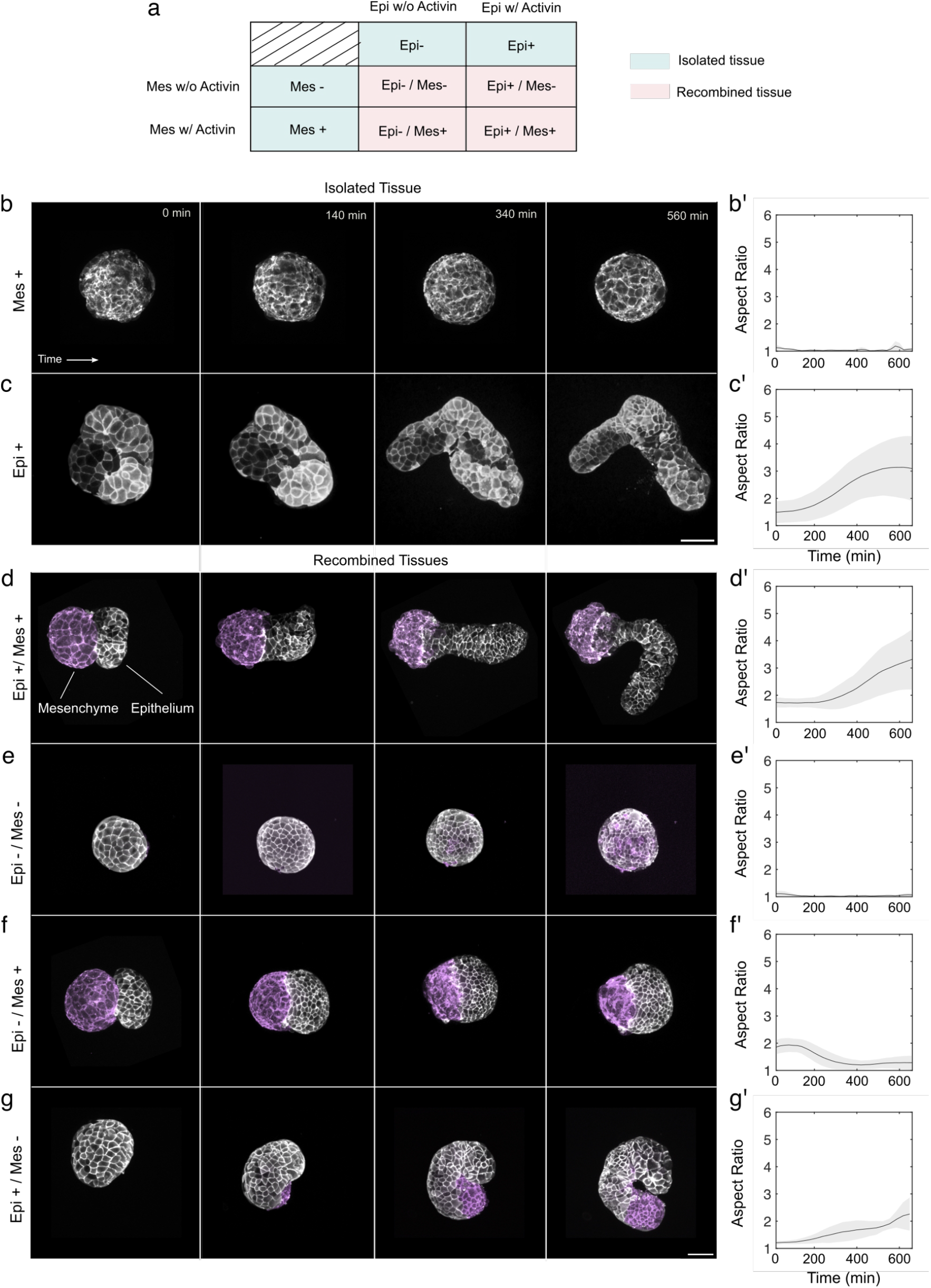
Elongation behaviors of isolated and recombined explants in different conditions. A) Table of isolation and recombined tissue experiments. Tissues are isolated with Activin or not and tested for elongation in isolation or recombined. This yields a total of eight conditions. B-C’) Isolated mesenchyme and epithelial tissue experiments and their respective aspect ratio plots. N=5 D-G’) Recombination experiments and their respective plots. N=5 for all conditions. Scale bar = 100µm

To further interrogate the roles of the two tissues, we recombined the isolated mesenchyme and epithelium, each stimulated or not with Activin. This procedure yields four possible combinations (Fig. 2A). As expected, if both tissues were Activin-stimulated (Epi +, Mes +), the recombined tissues elongated, albeit slightly less than explants that were kept intact (compare Fig. 2D, D’ to Fig. 1B,C). Meanwhile, explants recombining two unstimulated tissues (Epi -, Mes -) did not elongate (Fig. 2E, E’, Movie S5). When recombining a non-stimulated epithelium (Epi -) with a stimulated mesenchyme (Mes +), both tissues start slightly separated before the epithelium engulfed the mesenchymal tissue (Fig. 2F, Movie S5). In contrast, the explants recombining a stimulated epithelium (Epi +) with a non-stimulated mesenchyme (Mes -) elongated, but in a more disordered fashion and to a lesser extent than any of the fully stimulated explants (compare Fig. 2G, G’ to Fig. 2D, D’ & 1B, C). Thus, only recombined explants with an Activin-stimulated epithelium displayed any elongation.

Taken together, in contrast with previous works that have emphasized the crucial role of an active mesenchyme, our data show that an Activin-stimulated epithelium is required for elongation. While the Activin-stimulated epithelium, isolated or recombined, is capable of driving elongation, it does so more robustly and to a greater extent when combined with Activin - stimulated mesenchymal cells. When we fixed Epi +, Mes + cases we found that the epithelium would elongate without internal mesenchymal cells (Supplemental Fig. S3) and localisation of tissue extension is still constrained to the boundary in the absence of mesenchymal cells (Supplemental Fig. S4, Movie S6).

In the mesenchyme, two modes of active cell intercalation driving Xenopus axis elongation are known, anisotropic interface tension and crawling, are known to exist the mesenchymal cells (Weng et al., 2022). However, less is known about the epithelium. To identify which mechanism might drive epithelium-driven elongation, we carried out vertex simulations to find potential signatures of the dominant mechanism.

### Only in actively elongating tissues cells can orient perpendicular to the elongation axis

To better understand the signatures of different kinds of active tissue elongation, we carried out vertex model simulations (Fig. 3a). The vertex model describes biological tissues as polygonal tilings, where each cell corresponds to a polygon, and the passive response of a cell to an applied force is defined by the existence of both a preferred cell area and a preferred cell perimeter (details in the Methods). Using this model, we studied the behaviour of three different kinds of tissues (Fig. 3c-e): (i) a passive tissue and two tissues with different active CE mechanisms: (ii) active anisotropic cell-cell interface tensions and (iii) active crawling forces leading to cell-cell intercalation. We analyzed the behavior of these tissues for different imposed CE rates *V*, which we measure in terms of the natural relaxation time of the tissue to external stimuli. We observed the required external force to maintain this CE rate *V* (Fig. 3f) and a so-called nematic cell shape order parameter *Q* (Fig. 3g).

**Figure 3.**
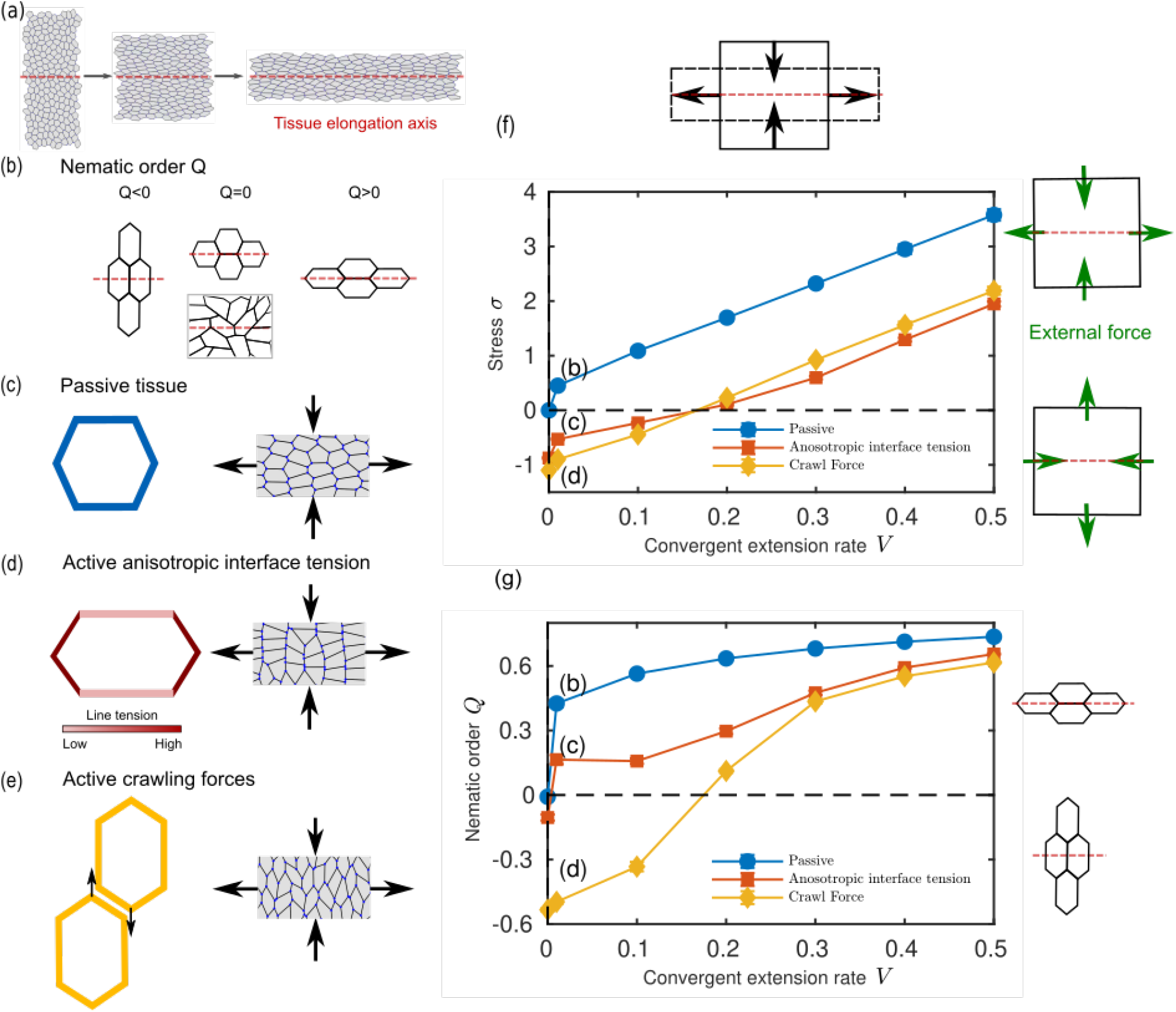
Computational model: a perpendicular cell shape alignment is only possible in actively deforming tissues. (a) We carry out vertex model simulations by progressively deforming a rectangular region of tissue, forcing it to undergo convergent-extension (CE) at a given rate *V* (axis of extension marked by a red dashed line). The vertex model describes each cell as a polygon with an area elasticity and cell-cell interfacial tensions (details in the Methods). (b) We describe cell shape in our simulations using a cell shape order parameter *Q* (definition in the Methods section), which describes the average cell shape anisotropy and its alignment with the axis of tissue elongation (red dashed line). Whenever cells are on average aligned parallel (perpendicular) to the axis of tissue extension, *Q* is positive, *Q*>0 (*Q* is negative, *Q*<0). When cell shapes are either isotropic or when they are anisotropic but oriented in random directions, *Q* is zero, *Q*=0. (c-e) We consider three different types of tissue dynamics, a purely passive tissue (c), a tissue with active anisotropic interface tensions (d), and a tissue where cells actively crawl past each other (e). (f) We find that the mechanical force *σ* required to achieve a certain CE rate *V* increases almost linearly with *V*, yet it is smaller for the active tissues (yellow and orange curves). For both kinds of activity, the activity is sufficient to drive CE at smaller rates *V* without any positive externally applied mechanical force *σ*. (g) For passive tissues and for tissues with line tension anisotropy, cell shapes are almost always aligned parallel to the axis of tissue extension. Yet, for anisotropic deformation driven by crawling forces, the cell shapes become perpendicular to the axis of extension, as long as the extension is not accelerated by externally applied forces.

The cell shape order parameter *Q* combines both; information about cell shape anisotropy and cell shape alignment with the axis of tissue elongation. Thus, if *Q*=0, cells are either not elongated, or they are individually elongated but not aligned in any way with the axis of explant elongation (Fig. 3b). A negative *Q* indicates that cells are both elongated and aligned perpendicular to the explant axis, and a positive *Q* indicates that cells are both elongated and aligned parallel to the explant axis (Fig. 3b).

To elongate the purely passive tissue (i), a positive external force in the elongation direction was always required, which increases with the elongation rate *V* (blue curve in Fig. 3f). Moreover, we found that *Q* was positive and large, even if the tissue is elongated very slowly (blue curve in Fig. 3g), indicating a strong alignment of cell shape parallel with the elongation axis.

When the tissue with an anisotropic interface tension (ii) was elongated with a CE rate *V* of approx. 0.2, no externally applied force was needed for elongation (orange curve in Fig. 3f). For a faster CE, external forces in the elongation direction were required, while for a slower CE, opposing external forces needed to be applied (red curve is positive/negative for larger/smaller *V* in Fig. 3f). In these simulations, cell shape was always parallel to the extension direction, indicated by a positive *Q* (red curve in Fig. 3g). Yet, when including fluctuations, we observed some small amount of cell shape alignment perpendicular to the direction of active extension (Supplemental Figure S5), consistent with recent work (Duclut et al., 2022). Taken together, this mechanism allows for autonomous CE even in the presence of opposing forces. Without fluctuations, this always led to a parallel alignment of cell shapes with the elongation axis.

When the tissue with crawling forces (iii) was elongated with a CE rate *V* of approx. 0.2, no externally applied force was needed (yellow curve in Fig. 3f), and an in- or decrease of the elongation rate required the presence of external forces, like for the tissue with anisotropic interface tension (ii). However, in the presence of crawling forces, cell shape could also align perpendicular to the elongation axis (negative value in yellow curve in Fig. 3g, compare Fig. 3e). This occurred for small CE rates *V*, which required opposing external forces (yellow curve in Fig. 3f). Meanwhile, for an elongating tissue without any external forces, our results suggest essentially isotropic cells (compare intersections of yellow curves in Figs with zero). Thus, we find a perpendicular cell shape alignment for the crawling force mechanism if the elongation is opposed by some external force.

Taken together, our vertex model simulations suggest that the cell shape elongation and alignment parameter *Q* indicates (i) whether tissue elongation is purely driven by externally applied forces or by internally generated active forces, and (ii) what kinds of mechanisms could create the internal forces. In particular, cells that align perpendicular to the direction of tissue elongation are a clear indicator for an internally driven tissue elongation, likely driven by cell crawling and opposed by some external forces. In the case of our Xenopus explants, such opposing forces could come for instance from an effective viscous friction of parts of the involved tissues.

### Cells in both mesenchymal and epithelial tissues align perpendicular to the axis of tissue elongation

To determine which mechanism drives cell intercalation in our explants, we quantified the cell shape order parameter Q. In intact explants, epithelial cells gradually orient perpendicular to the axis of elongation (Fig. 4A, top panel), suggesting that the epithelium contributes to explant elongation. Moreover, individual movies show that in the region close to the mesenchymal-epithelial interface elongating most strongly, the cell orientation perpendicular to the local explant axis is the most pronounced (Fig 4B, top, black arrowheads and Movie S7).

**Figure 4.**
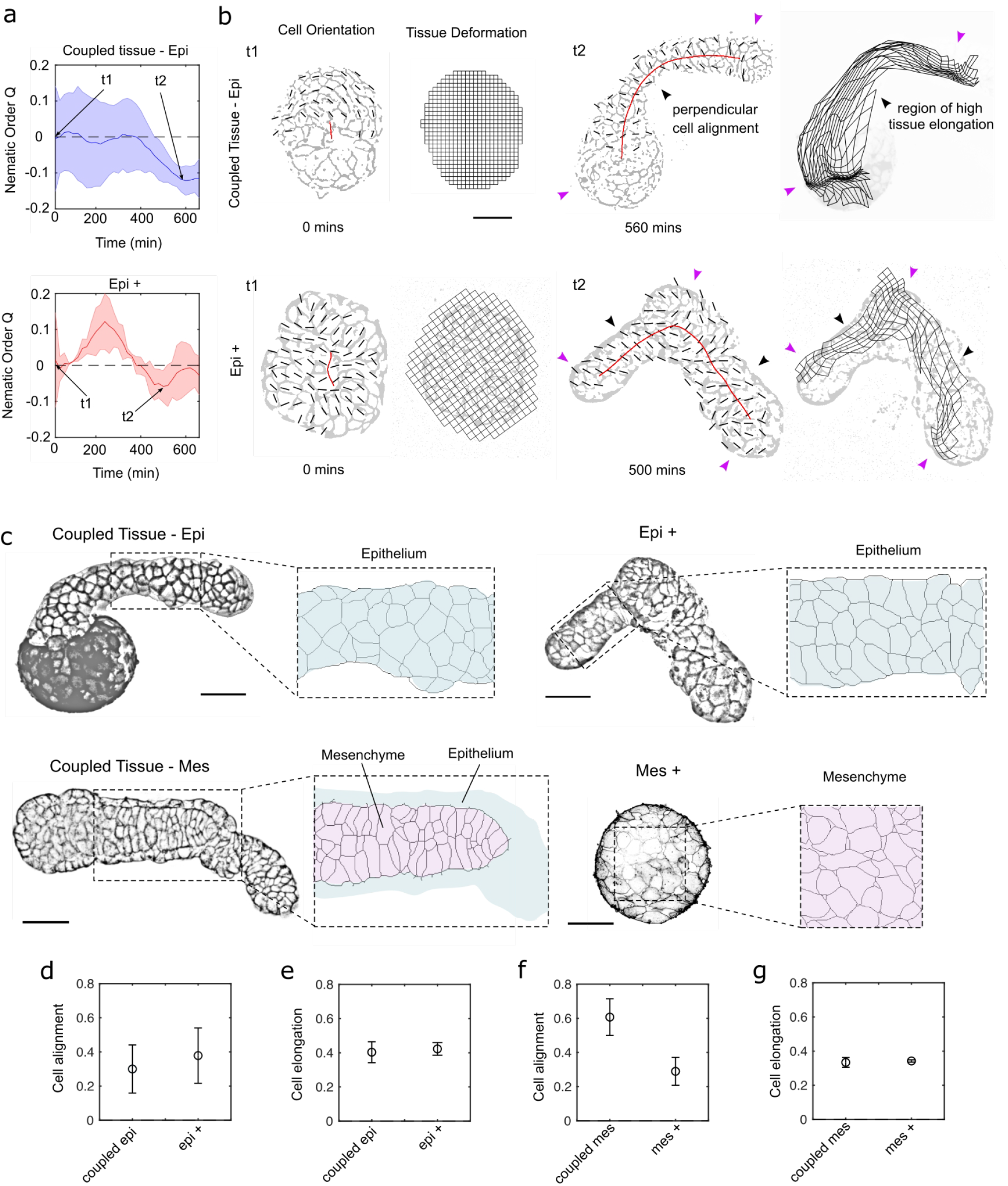
Cell shape elongation and alignment. A) Plots of average nematic order Q of epithelial tissue over time in (N=4) for intact (top) and isolated conditions (bottom). B) Snapshots of explants showing the local nematic order and epithelial tissue deformation for intact (top) and isolated (bottom) explants at early (t1) and late (t2) timepoints. Black arrows indicate regions that show high anisotropic deformation; magenta arrowheads show low anisotropic deformation. C) Tissue organization in coupled and isolated tissues for both epithelium and mesenchyme. (N=3) D) Comparison of decomposed nematic order Q (cell alignment and cell orientation) in intact coupled tissue epithelium and mesenchyme and in isolated epithelium and mesenchyme. Scale bar = 100µm for all

In the isolated Epi+ explants, we observed more variability in cell orientation and elongation when averaged over multiple explants (Fig. 4A, bottom). Yet individual movies again show that regions that locally elongate more strongly display more frequently a cell orientation perpendicular to the local explant elongation axis (Fig 4B bottom, black vs. magenta arrowheads and Movie S7).

Taken together, these data show that the epithelial cells orient and order over time perpendicular to the elongation axis, coupled to or in the absence of the mesenchyme. According to the model, this supports the idea that epithelial cells actively drive elongation by a crawling mechanism. We also studied cell shapes in the mesenchyme. Yet, because Xenopus tissues are opaque, we needed to base our analysis on fixed explants. Strikingly, we found that in intact explants, mesenchymal cells also show a strong alignment perpendicular to the explant axis (Fig. 4C). In isolated Mes+ explants, individual cells are also elongated, but cells on the whole are less aligned. This suggests that mesenchymal cells also contribute to actively drive explant elongation, yet their alignment and orientation depends on the presence of an epithelium.

### Perpendicular cell orientation first appears in the basal epithelium

To better understand the interactions between epithelium and mesenchyme in defining cell shape alignment, we imaged both epithelial and mesenchymal tissues from the most external surface of the epithelium to the deep mesenchymal cells and quantified the cell shape parameter Q (Fig. 5A). We study cell shape alignment at different time points, including times when the explants are not yet elongating. Thus, we take as a spatial landmark the boundary between mesenchymal and epithelial tissues instead of the explant elongation axis (Fig. 5A).

**Figure 5.**
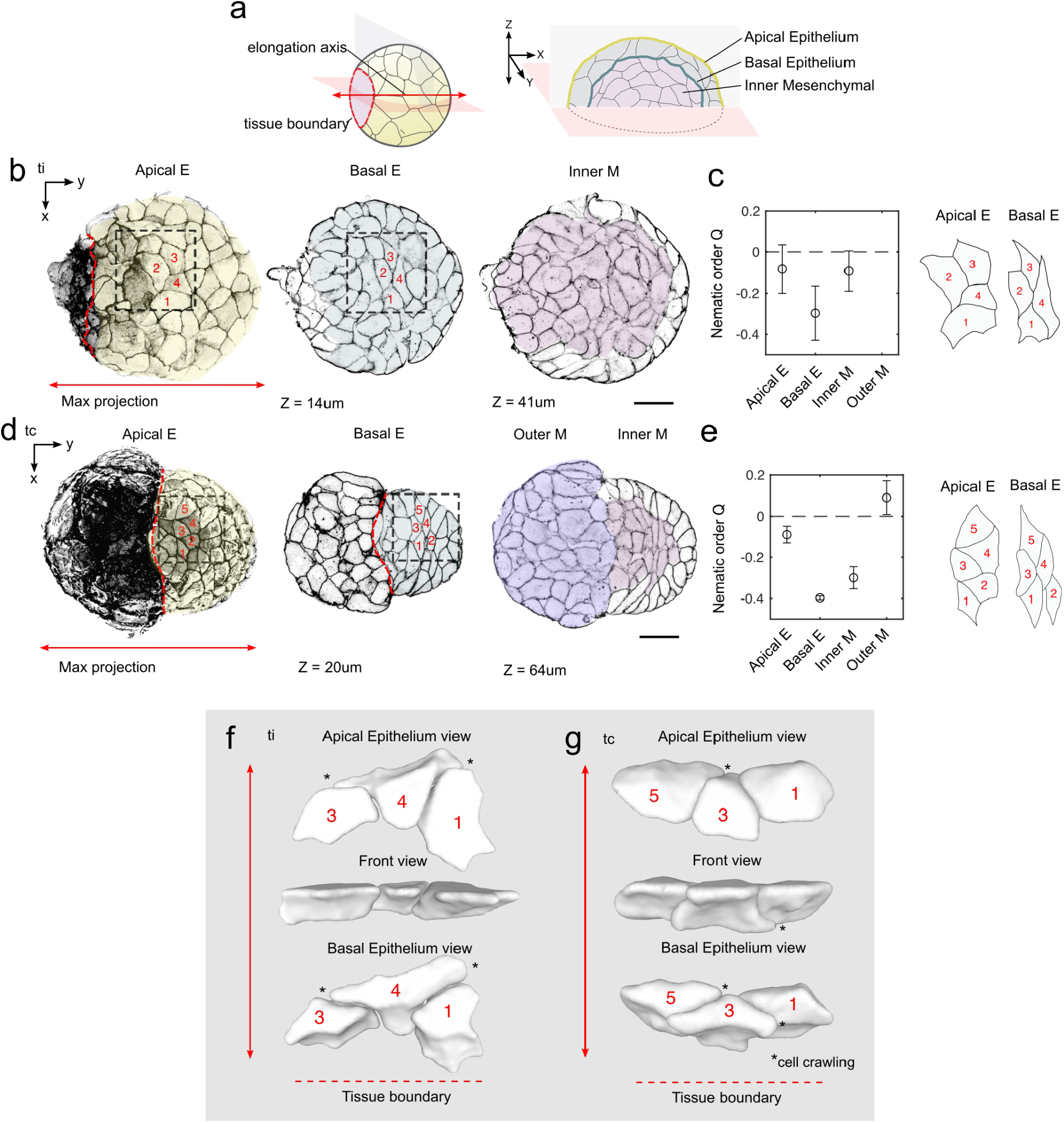
Emergence of cellular orientation parallel to the tissue boundary originates in the basal epithelium and propagates to the internal mesenchymal cells. A) Diagrams showing explant boundary and predicted axis of elongation and sectioned explant showing apical epithelium, basal epithelium and first layer of inner mesenchymal cells in the Z axis. B) Images and segmentations with cell orientation representation for three Z positions at timepoint ti C) Representative close-up of segmented cells and corresponding nematic order Q score for each Z position (N=3). No outer mesenchymal cells are quantified at this time point as mesenchymal cells are predominantly covered by epithelium. D) Images and segmentations with cell orientation representation for three Z positions and at time point 2. F) Representative close-up of segmented cells and corresponding nematic order Q score for each Z position (N=3). F) 3D segmentation of epithelial cells at initial timepoint tc showing differences in cellular elongation and orientation between the apical and basal surface. * shows where basal epithelium shows signs of basal crawling behavior. G) 3D segmentation of epithelial cells at time point tc showing differences in cellular elongation and orientation between the apical and basal surface. * shows where basal epithelium shows signs of basal crawling behavior.

We found that in early explants that are still rounded up, the basal cell cross-sections of the epithelium were strongly aligned parallel to the tissue boundary (=perpendicular to the later elongation axis, Fig. 5B, C). Meanwhile, the apical cell cross-sections of the epithelium and the mesenchymal cells were more disordered (Fig. 5 B, C, Movie S8).

At later times when the epithelium has contracted, but before explant elongation, the basal cell cross-sections of the epithelium were even more sharply aligned parallel to the tissue boundary as compared to the apical cross-sections (Fig. 5D, E, Movie S8). Interestingly, now mesenchymal cells are also highly aligned parallel to the tissue boundary (Fig. 5D, E, Movie S8). Meanwhile, mesenchymal cells that are now squeezed out of the explant do not show any clear alignment (Fig. 5D, E, Movie S8).

Taken together, our data indicate that: (i) the epithelial basal surface is the origin of oriented bipolar activity which is always oriented parallel to the tissue boundary, (ii) there is a transfer of orientation order from the basal epithelial cells to the underlying mesenchymal cells, and (iii) the emergence of basal epithelial and mesenchymal cell shape order precedes that of the apical epithelium and explant elongation

Given that the epithelial cells always align parallel to the tissue boundary, we wondered whether the tissue boundary is responsible for setting their orientation, and subsequently the elongation axis. To test this, we split explants into either two or three pieces and recombined them (Fig 6, A and B). The mesenchyme tissues of the recombined explants fused, resulting in single aggregates. However, two or three boundaries were created at the edges of the covering epithelial pieces, as seen in Fig 6A (time 0 min). The round aggregates underwent dramatic shape changes within a few hours. The two-piece aggregates acquired an elongated shape, with two symmetric extensions away from the parallel boundaries (Fig 6C, Movie S9). The three-part aggregates acquired a triangular shape, the vertices of which lie in a direction bisecting the initial boundaries (Fig 6B, Sup Mov). These observations suggest that the tissue boundaries set the elongation axes and, therefore, the final shape of the aggregates. This shows that the initial conditions of the tissue coupling provide direction cues for morphogenesis.

**Figure 6.**
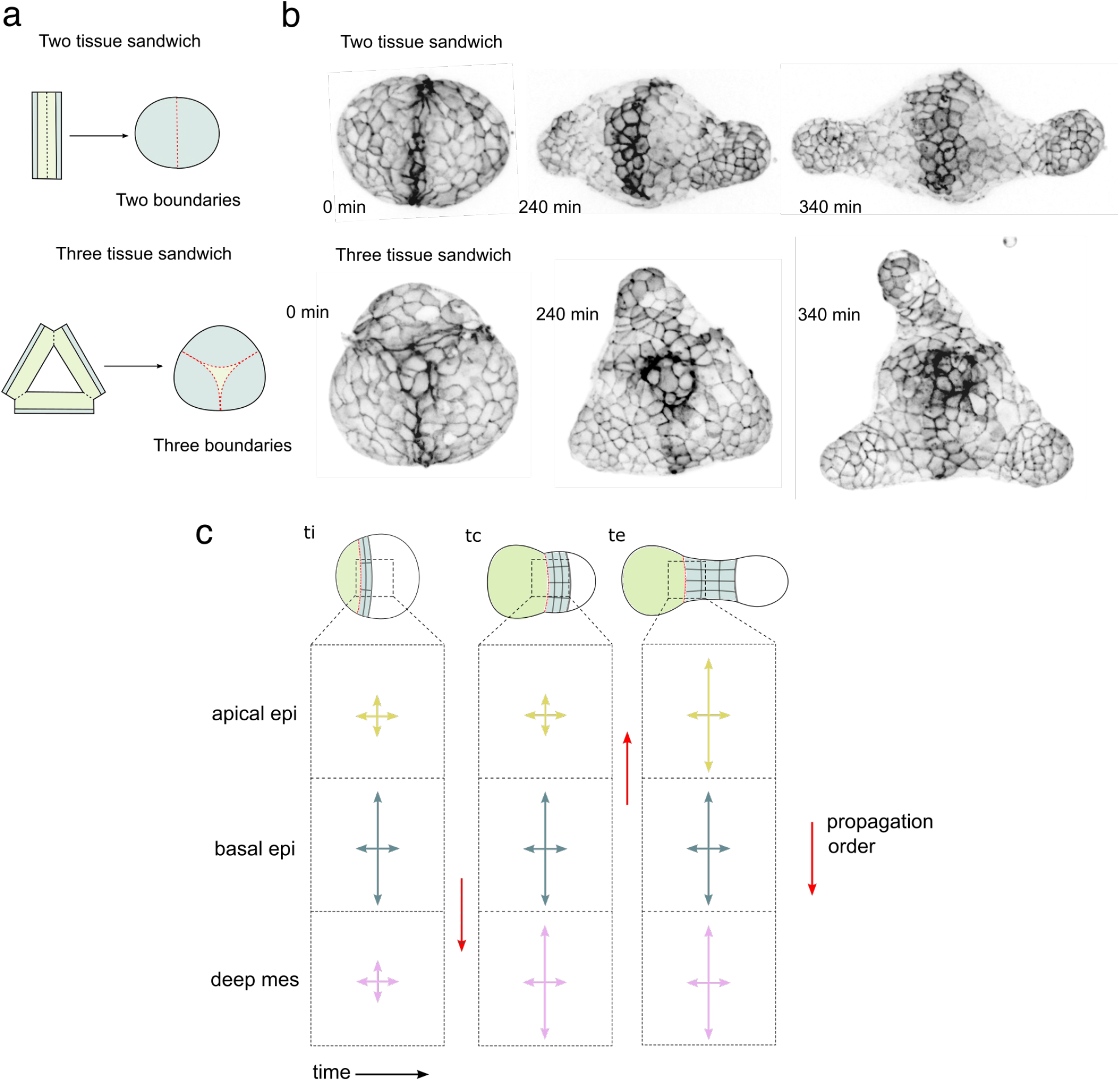
Additional boundaries create additional elongation poles. A) Schematic of two and three tissue experiments B) Snapshots of two and three tissue experiments over time C) Schematic of cellular alignment propagation at progressive timepoints

## Discussion

In this paper, we use Xenopus animal cap explants to uncover how tissues self-organise to establish and extend an axis in the absence of external positional information or directional cues (Figure 1). Previous work on axis extension focussed on mesenchyme alone, studying for instance, polarised cell-cell interactions (Davidson et al., 2006) (Skoglund et al., 2008) (Shindo and Wallingford, 2014). Here, we complement these studies by uncovering the importance of the mesenchymal-epithelial interplay. We show that both mesenchymal and epithelial tissues drive axis elongation but in a hierarchical fashion. The epithelial tissue can elongate by itself, while the mesenchymal tissue requires the initial presence of the epithelium in order to elongate. This is likely because explant elongation requires aligned and elongated cells. Such alignment is visible initially in the epithelium, from which it propagates to the mesenchyme. Our results suggest that this cellular alignment, and thus the axis of explant extension, is set by tissue boundaries, e.g. between the epithelium and mesenchyme.

Our live imaging and tissue isolation and combination experiments reveal that only when an Activin-stimulated epithelium is present, explants will elongate or polarise underlying mesenchymal cells. Indeed, previous work has shown the epithelium to have organiser-like effects; when grafts of epithelium from the dorsal marginal zone are transplanted to the ventral side of host embryos, they induce a second axis (Shih and Keller, 1992). However, these findings seem to contradict a previous report that suggested the role of the epithelium was passive (Ninomiya and Winklbauer, 2008), because explant extension did not depend very strongly on the kind of epithelium used. Yet, in their experiment, the mesenchymal cells came from the lip, where they are likely already polarized, e.g. in terms of cell shape or PCP. Meanwhile, by virtue of using our early stage-9 animal cap explants, which are in a more naive and unpolarized state, we find that mesenchymal cells from the animal cap are disordered and require an Activin-stimulated epithelium to align them. We find that such cell shape alignment proceeds in a 3-step mechanism across tissues: (1) before the onset of elongation the basal surfaces of epithelial cells elongate and orient parallel to the epithelium-mesenchyme boundary; (2) the inner mesenchymal cells align in the same direction; (3) the apical surfaces of epithelial cells orient parallel to their basal surface (Fig 6).

We provide several pieces of evidence that the epithelial-mesenchyme (epi-mes) boundary defines the direction of cell shape alignment and explant elongation. First, both cell alignment and CE occur only in a narrow band close to the boundary where the epithelium covers the mesenchyme (Fig 1 and Fig 5). Second, an isolated Activin-stimulated epithelium, which lacked any epi-mes boundary, did not undergo consistent extension. Third, when multiple ectopic boundaries were created, several corresponding elongation axes formed (Fig. 6). That the tissue boundary can provide directional cues has been suggested before (Green et al., 2004), where Activin-stimulated dissociated mesenchymal cells were made to aggregate into a flat disk before rounding up into a ball and elongating perpendicular to the original disk. While in those experiments the boundary was defined by the experimenter, it emerged on its own in our experiments. Together, this shows that tissue boundaries set the conditions for coherent axis elongation.

Our work raises several intriguing questions for future research. First, it will be interesting to dissect the propagation mechanism from the boundary to the inner cells and along the axis of elongation. Further evidence of epithelial tissue signaling shows perpendicular unidirectional microtubule growth in chordamesoderm cells adjacent to ectodermal tissue that is prior to shape elongation (Shindo et al., 2008). Interestingly, this was found to be independent of PCP signaling. Second, while we showed that the mesenchymal cells align with the overlying epithelial cells, sample scattering did not allow us to specify the full 3D shape of the mesenchymal cells over time.

Third, our results give some indications as for the nature of the cellular forces driving convergence-extension. Generally, oriented cells may drive convergence-extension through cell crawling and/or apical junction shrinkage (ref). Our simulation results suggest that perpendicular alignment of cell shape to the elongation axis occurs more easily for a cell crawling mechanism. Furthermore, we also observed protrusions in the basal part of the epithelial cells, where such perpendicular alignment first occurs. This suggests a cell crawling mechanism, but future work is required to ascertain this point.

Fourth, our animal cap explants share similarities with the Xenopus exogastrula, where the embryo is made to gastrulate externally (Tucker and Slack, 1995). In particular, such exogastrulas elongate exclusively in a region close to the boundary between the mesoderm and ectoderm. More generally, boundary creation could provide a generic mechanism for cells to reliably align in the same manner, thereby guiding morphogenesis and suppressing noise.

## Materials and Methods

### Xenopus Laevis

All experiments were performed following the Directive 2010/63/EU of the European parliament and of the council of 22 September, 2010 on the protection of animals used for scientific purposes and approved by the “Direction départementale de la Protection des Populations, Pôle Alimentation, Santé Animale, Environnement, des Bouches du Rhône” (agreement number G 13055 21) as (Walton et al., 2024).

### Embryo Culture

Ovulation was stimulated in *X. laevis* adult females by injection of (800 units/animal) Human Chorionic Gonadotropin (ChorulonR). On the following day, eggs were recovered by squeezing, fertilized *in vitro* with sperm from Nasco males, de-jellied in 2% cysteine hydrochloride (pH 8.0) and washed, first in water, then in 0.1X MBS (Modified Barth’s Saline). Embryos were kept at 13°C or 18°C until ready for injecting at 2-4 cells. They were then transferred to 4% Ficoll in 1X MBS for at least two hours for injection puncture healing. Injections were done using pulled glass capillaries and embryos were injected either twice in each blastomere at the 2-cell stage or once in each cell at the 4-cell stage with 20nl total volume of mRNA solution to target the whole animal cap.

### Animal cap explants

Animal caps were excised at stage 8.5-9.5, washed once in 1X MBS and transferred to Lipidure coated plastic dishes with 5nl/mL of Activin A (R&D systems) in 1X MBS. Each cap that was excised was cut using an eyelash knife into two equally sized smaller squares. For isolated and recombined tissue experiments the animal cap outer epithelium was excised directly from the whole embryo using an eyelash knife and forceps and the underlying deeper mesenchymal cells were dissected out afterwards before being separated out in another dish. Separate mesenchymal or epithelial tissues were soaked with Activin A and recombined after two hours in a new dish with just 1X MBS. Care was taken to recombine tissues by gently pushing them together just before the onset of gastrulation when they are less responsive to Activin A.

### mRNA Synthesis

GPI-GFP and Membrane-RFP plasmids were kindly provided by Laurent Kodjabachian. Sense capped mRNAs were synthesized from linearized plasmids with the Sp6 Ambion mMessage mMachine kit (Life Technologies), and purified with Macherey-Nagel NucleoSpin RNA Clean-up kit as (Walton et al., 2024). After determination of the concentration, aliquots were kept at - 80C. GPI-GFP was injected at 500 pg per cell at 2 or 4 cells while Membrane-RFP was injected at 250 pg per cell at 2 or 4 cells.

### Spinning disk and Confocal Imaging

To avoid rotation of explants during live imaging, they were transferred to glass bottom dishes (Mattek) coated with lipidure (Lipidure) and filled with 1.8% methylcellulose (Sigma) in 1X MBS. Explants were gently pushed down to the glass bottom for live imaging. Time-lapse imaging was done at 20°C on a spinning disk microscope (Nikon Eclipse Ti) using a 10X air objective. Image acquisition was acquired with metamorph software. Explants were imaged 100µm deep with 5µm steps every 20 mins for 14 hours.

Fixed explants were embedded in RapiClear 1.52 (SunJin Lab) for optical clearing and sandwiched between two coverslips with 120 µm spacers. Explants were then imaged using an inverted confocal microscope (Zeiss LSM 880) with a 60X objective at 1µm Z steps.

### Measuring aspect ratio of curved, irregular shaped explants

To measure aspect ratio, Max-projected time lapses were registered using the ImageJ Plugin MultiStackReg to remove translational movements. Masks were created using Gaussian blurring (sigma 2) and thresholding in ImageJ. A custom made Matlab code was used to create distance maps and skeletonised images of the masks where the elongation axis were obtained by obtaining the length of the points in the skeletonised image, plus the two end points of the skeleton on the distance map. For the width an average of all the widths along the elongation axis was taken using the distance map.

### Tissue deformation tracking

To track tissue deformation we used particle image velocimetry (PIVlab) of explant time lapses, using 3 decreasing box sizes, where the last box size is larger than a cell size. The masks are used to constrain the PIV analysis to the explant region. A custom-made Matlab code was used to create a grid confined to the explant that deforms when advected at each time point using vectors obtained from PIV analysis.

### Cell interface alignment with elongation axis

To track cell interface alignment we processed time-lapse images as follows. i) a duplicate stack of the time-lapse was made with added Gaussian blur (sigma 10) in ImageJ ii) the blurred image was divided by the original image using image calculator iii) then the 32 bit image was thresholded until the best signal to noise was reached by observation. We used a custom-made Python script (described in detail below) to compute nematic order Q. The nematics were fed into a custom Matlab code which compared each nematic orientation to the closest local tangent on the midline axis (obtained from skeletonised masks) and compared the two orientations to compute how cell interfaces align with the axis. Cell interface alignment was also computed in fixed samples that were segmented using Cellpose (Fig. 4).

### 3D segmentation and visualization of polarized epithelial cells

3D segmentation was done using LimSeg which creates a triangular mesh that grows from a spherical seed until it reaches the outer boundaries of the selected cells in a stack. We used a D_0 =3.9 which is a parameter that defines how fine the detail you want to capture and F_pressue=0.015 which is related to the expansion force of the active contour of the seeded sphere. We then visualized the meshes, adding shading and bone color using the application MeshLab 2023.

### Computation of the cell shape order parameter *Q*

In the simulations, where the axis of tissue extension is the *x* axis, we define the cell shape order parameter *Q* as follows:

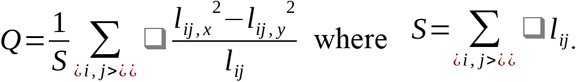

Here, both sums are over all cell-cell interfaces *ij*. The length vector of interface *ij* is denoted by (*l*_*ij,x*_, *l*_*ij,y*_), and its length is *l*_*ij*_.

To compute *Q* from an experimental image, we first quantify the local orientation of cell interfaces using the approach one of use developed earlier to quantify Drosophila wing hair orientation from (Sagner et al., 2012). We modified it for visualization by adjusting the box size to fit approximately one cell. For each box *b*, we then determined the closest place on the midline (defined as described above in section Image Analysis), and took the local tangent of the midline as a reference direction whose angle we denote by *θ*_*b*_. We then compute *Q* as:

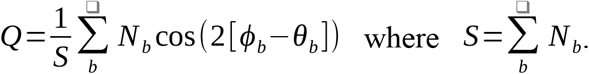

Here, both sums are over all boxes *b*. In a given box the nematic magnitude is denoted by *N*_*b*_, and the angle of the nematic is denoted by *ϕ*_*b*_.

Note that in the limit of straight cell-cell interfaces, straight midline, high image resolution, and small box size, both simulation and experimental definitions of *Q* exactly match.

### Model

We simulate the tissue with *N*_*cells*_ cells using a 2D vertex model, which describes a tissue by a packing of polygons, each of which represents a cell in the tissue. We call the polygon corners vertices. The tissue is subject to to periodic boundary conditions, where the periodic box size can change over time, while the total area remains constant *L*_*x*_*L*_*y*_ = *N*_cells_*A*_0_, where *A*_0_ is a constant parameter denoting the preferred cell area (see next paragraph).

To define the mechanical behavior of the system, we introduce the following mechanical work function:

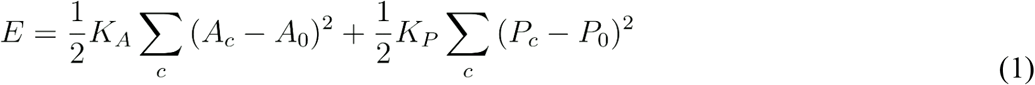

Both sums in this equation are over all cells *c* of the tissue. The first term represents an area elasticity of cells, where *A*_*c*_ is the area of cell *c*, while *A*_0_ and *K*_*A*_ are constant parameters: *A*_0_ is the preferred cell area, and *K*_*A*_ is the associated area stiffness. The second term describes a cell perimeter elasticity, where *P*_*c*_ is the area of cell *c*, while *P*_0_ and *K*_*p*_ are constant parameters: *P*_0_ is the preferred cell perimeter and *K*_*p*_ is the associated perimeter stiffness. The work function *E* defines elastic forces on the individual vertices, where the elastic force on some vertex *i* is given by the partial derivative *∂E/∂r*_*iα*_, where *r*_*iα*_ denotes the position of vertex *i*. Here and below, Latin letters *i, j*,… denote vertex indices and Greek letters *α, β*,…ϵ{*x,y*} denote dimension indices.

We impose a constant rate *V* of pure shear deformation of the periodic box along the *x* axis. In other words, the periodic box dimensions follow the following dynamics with time *t*:

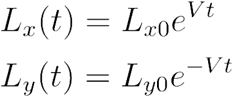

Here, *L*_0*x*_ × *L*_0*y*_ are the initial dimensions of the periodic box.

To describe the time evolution of the vertex positions, we take into account friction terms on different geometric elements, such as cell area, cell perimeter and interface length. We also include a small substrate friction to regularize the dynamics. The instantaneous vertex velocities 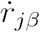 are extracted from solving the following linear problem:

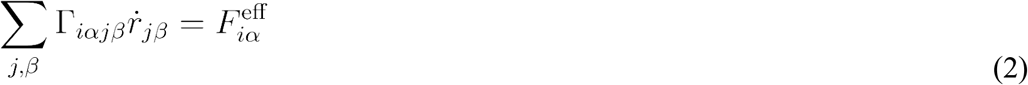

Here Γ_*iαjβ*_ is a friction matrix and 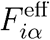 is an effective force, respectively defined as

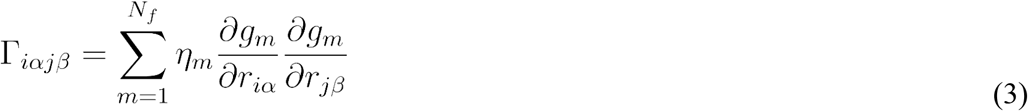

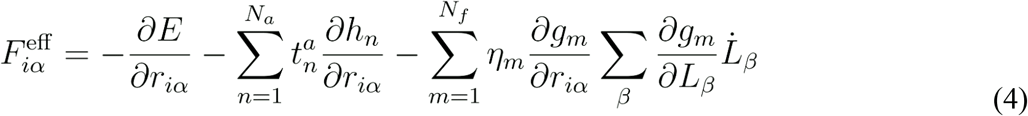

In both equations, *N*_*f*_ is the total number of geometric elements *m* on which friction acts, each associated with a friction coefficient *η*_*m*_. Each geometric element *m* is a scalar quantity *g*_*m*_ (cell area, cell perimeter, interface length, and vertex *x* and *y* positions). For instance, the first 2*N*_cells_ of the *N*_*f*_ frictional elements *g*_*m*_ are given by cell areas and perimeters; i.e., with the cells labeled *c* = 1,…,*N*_cells_, we have:

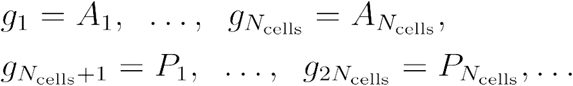

The remaining frictional elements *g*_*m*_ are set to all interface lengths and the *x* and *y* positions of all vertices.

In Eq. (4) the first term comes from the elastic interactions among the vertices. The second term is a contribution from *n* = 1,…,*N*_a_ active tensions 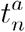 acting on geometric elements *h*_*n*_. The active tensions are implemented differently for different kinds of tissues (passive, active anisotropic interface tensions, active crawling). We present the concrete implementation of this term in the next paragraph. The third term in Eq. (4) is due to deformations of the periodic box.

We simulate the vertex model for three different tissues: (i) passive tissue, (ii) active tissue with anisotropic interface tensions, and (iii) active tissue with cell crawling forces :

i. To simulate passive tissues we do not include any active tensions 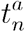, i.,e. we set *N*_a_ = 0 in Eq. (4).
ii. To simulate tissues with an active, anisotropic interface tension, we include an active tension 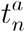 for each cell-cell interface *ij*. Thus, the active tension index corresponds to the interface index (composed of two vertex indices), *n* ≡ *ij*, and the geometric elements are the cell-cell interface lengths, *h*_*n*_ ≡ *h*_*ij*_ ≡ *h*_*ij*_. For each interface *ij* we define an active tension 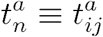 that is modulated by the instantaneous angle *θ* of the interface with respect to the *x* axis. Additionally, in order to compare to previous results in the literature (Duclut et al., 2022), we also include Ornsetin-Uhlenbeck noise *μ*_*ij*_ into the active tension:

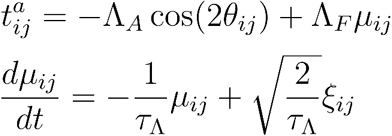

Here, Λ_*A*_ denotes the interface tension anisotropy amplitude; this part of the active interface tension is −Λ_*A*_ for interfaces oriented along the *x* axis and +Λ_*A*_ for interfaces oriented along the *y* axis. +_*F*_ denotes the fluctuation amplitude, *τ*_Λ_ is the characteristic Ornstein-Uhlenbeck time scale, and *ξ*_*ij*_ are zero-mean, unit-variance, independent Gaussian white noise sources.
iii. To simulate tissues with active crawl forces, we introduce for each cell *c* pushing forces with constant magnitude *F*_*a*_ that the vertices with minimal and maximal *y* positions exert on their neighbors. Yet this alone would violate linear and angular momentum balance (remember that we describe a tissue that is not linked to a solid substrate). To compensate for this, we apply counteracting forces of appropriate magnitude on the two neighboring cell centers (Figure S6).

Within the formalism of Eqs. (2)-(4), linear and angular momentum are automatically conserved when using scalar elements *g*_*m*_ and *h*_*n*_. In this formalism, the described active crawling forces correspond exactly to defining *N*_a_ = 2*N*_cells_ geometric elements with active tensions in the following way. We pick within each cell *c* the two vertices *i* and *j* with minimal and maximal y position, respectively, and we define a geometric element for each of them: *n* ≡ *ci*. For vertex *i*, we consider the intersection between on the one hand the cell-cell interface next to vertex *i* that does not abut cell *c* and on the other hand the connection line between the barycenters of the two neighbor cells of c that abut vertex *i* (see Figure S6). We define the geometric element *h*_*n*_ ≡ *h*_*ci*_ as the distance between this intersection and vertex *i*. The corresponding active tension is then set to a constant parameter 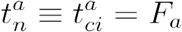. We proceed exactly analogously for vertex *j*.

We solve Eqs. (1)-(4) using an Euler scheme with an adaptive time step that ensures that the error in the vertex velocities is smaller than 10%. Equation (2) is inverted using a sparse matrix inversion through LDL^T^ Cholesky factorization, using the SimplicialLDLT class template from the C++ Eigen library.

### Simulation parameters

- Elastic parameters: *A*_0_ = 1, *K*_*A*_ = 10, *P*_0_ = 3.7, and *K*_*p*_ = 1.
- Frictions: *η*_*m*_ = 1 for all cell areas, cell perimeters, and interface lengths, and *η*_*m*_ = 10^−3^ for the vertex positions (i.e. substrate friction).
- The cut-off length for T1 transitions is 10^−1^.
- Active anisotropic line tension: amplitude Λ_*A*_ = 1. In the main text figure 3, orange curve, we did not include any fluctuations, Λ_*F*_ = 0. In Figure S5, the orange curve is the same as in the main text figure, while the … curve includes fluctuations with Λ_*F*_ = 0.5 and *τ* _Λ_ = 1.
- Crawling forces: The magnitude of the crawling forces is fixed to *F*_*a*_ = 1.

## References

Ariizumi, T., Takahashi, S., Chan, T., Ito, Y., Michiue, T., Asashima, M., 2009. Isolation and Differentiation of Xenopus Animal Cap Cells. CP Stem Cell Biology 9. 10.1002/9780470151808.sc01d05s9

Beloussov, L.V., Louchinskaia, N.N., Stein, A.A., 2000. Tension-dependent collective cell movements in the early gastrula ectoderm of Xenopus laevis embryos. Development Genes and Evolution 210, 92–104. 10.1007/s004270050015

Butler, L.C., Blanchard, G.B., Kabla, A.J., Lawrence, N.J., Welchman, D.P., Mahadevan, L., Adams, R.J., Sanson, B., 2009. Cell shape changes indicate a role for extrinsic tensile forces in Drosophila germ-band extension. Nat Cell Biol 11, 859–864. 10.1038/ncb1894

Christodoulou, N., Skourides, P.A., 2022. Distinct spatiotemporal contribution of morphogenetic events and mechanical tissue coupling during Xenopus neural tube closure. Development 149, dev200358. 10.1242/dev.200358

Collinet, C., Rauzi, M., Lenne, P.-F., Lecuit, T., 2015. Local and tissue-scale forces drive oriented junction growth during tissue extension. Nat Cell Biol 17, 1247–1258. 10.1038/ncb3226

Davidson, L.A., Marsden, M., Keller, R., DeSimone, D.W., 2006. Integrin α5β1 and Fibronectin Regulate Polarized Cell Protrusions Required for Xenopus Convergence and Extension. Current Biology 16, 833–844. 10.1016/j.cub.2006.03.038

Duclut, C., Paijmans, J., Inamdar, M.M., Modes, C.D., Jülicher, F., 2022. Active T1 transitions in cellular networks. Eur. Phys. J. E 45, 29. 10.1140/epje/s10189-022-00175-5

Green, J.B.A., Dominguez, I., Davidson, L.A., 2004. Self-organization of vertebrate mesoderm based on simple boundary conditions. Developmental Dynamics 231, 576–581. 10.1002/dvdy.20163

Holtfreter, J., 1944. A study of the mechanics of gastrulation. J. Exp. Zool. 95, 171–212. 10.1002/jez.1400950203

Keller, R., Davidson, L., Edlund, A., Elul, T., Ezin, M., Shook, D., Skoglund, P., 2000. Mechanisms of convergence and extension by cell intercalation. Phil. Trans. R. Soc. Lond. B 355, 897–922. 10.1098/rstb.2000.0626

Keller, R., Shih, J., Domingo, C., 1992. The patterning and functioning of protrusive activity during convergence and extension of the Xenopus organiser. Development 116, 81–91. 10.1242/dev.116.Supplement.81

Keller, R., Shook, D., Skoglund, P., 2008. The forces that shape embryos: physical aspects of convergent extension by cell intercalation. Phys. Biol. 5, 015007. 10.1088/1478-3975/5/1/015007

Kim, Y., Hazar, M., Vijayraghavan, D.S., Song, J., Jackson, T.R., Joshi, S.D., Messner, W.C., Davidson, L.A., LeDuc, P.R., 2014. Mechanochemical actuators of embryonic epithelial contractility. Proc. Natl. Acad. Sci. U.S.A. 111, 14366–14371. 10.1073/pnas.1405209111

Mancini, P., Ossipova, O., Sokol, S.Y., 2021. The dorsal blastopore lip is a source of signals inducing planar cell polarity in the Xenopus neural plate. Biology Open 10, bio058761. 10.1242/bio.058761

Maroudas-Sacks, Y., Garion, L., Shani-Zerbib, L., Livshits, A., Braun, E., Keren, K., 2021. Topological defects in the nematic order of actin fibres as organization centres of Hydra morphogenesis. Nat. Phys. 17, 251–259. 10.1038/s41567-020-01083-1

Munro, E.M., Odell, G.M., 2002. Polarized basolateral cell motility underlies invagination and convergent extension of the ascidian notochord. Development 129, 13–24. 10.1242/dev.129.1.13

Ninomiya, H., Elinson, R.P., Winklbauer, R., 2004. Antero-posterior tissue polarity links mesoderm convergent extension to axial patterning. Nature 430, 364–367. 10.1038/nature02620

Ninomiya, H., Winklbauer, R., 2008. Epithelial coating controls mesenchymal shape change through tissue-positioning effects and reduction of surface-minimizing tension. Nat Cell Biol 10, 61–69. 10.1038/ncb1669

Sagner, A., Merkel, M., Aigouy, B., Gaebel, J., Brankatschk, M., Jülicher, F., Eaton, S., 2012. Establishment of Global Patterns of Planar Polarity during Growth of the Drosophila Wing Epithelium. Current Biology 22, 1296–1301. 10.1016/j.cub.2012.04.066

Savin, T., n.d. On the growth and form of the gut.

Sawamura, K.-I., Uchiyama, H., n.d. Dose and time-dependent mesoderm induction and outgrowth formation by activin A in Xenopus laevis.

Shih, J., Keller, R., 1992. The epithelium of the dorsal marginal zone of Xenopus has organizer properties. Development 116, 887–899. 10.1242/dev.116.4.887

Shindo, A., Wallingford, J.B., 2014. PCP and Septins Compartmentalize Cortical Actomyosin to Direct Collective Cell Movement. Science 343, 649–652. 10.1126/science.1243126

Shindo, A., Yamamoto, T.S., Ueno, N., 2008. Coordination of Cell Polarity during Xenopus Gastrulation. PLoS ONE 3, e1600. 10.1371/journal.pone.0001600

Skoglund, P., Rolo, A., Chen, X., Gumbiner, B.M., Keller, R., 2008. Convergence and extension at gastrulation require a myosin IIB-dependent cortical actin network. Development 135, 2435–2444. 10.1242/dev.014704

Symes, K., Smith, J.C., 1987. Gastrulation movements provide an early marker of mesoderm induction in Xenopus laevis. Development 101, 339–349. 10.1242/dev.101.2.339

Tallinen, T., Chung, J.Y., Rousseau, F., Girard, N., Lefèvre, J., Mahadevan, L., 2016. On the growth and form of cortical convolutions. Nature Phys 12, 588–593. 10.1038/nphys3632

Tucker, A.S., Slack, J.M.W., 1995. Tail bud determination in the vertebrate embryo. Current Biology 5, 807–813. 10.1016/S0960-9822(95)00158-8

Walton, A., Thomé, V., Revinski, D., Marchetto, S., Puvirajesinghe, T.M., Audebert, S., Camoin, L., Bailly, E., Kodjabachian, L., Borg, J.-P., 2024. A vertebrate Vangl2 translational variant required for planar cell polarity. Journal of Biological Chemistry 300, 106792. 10.1016/j.jbc.2024.106792

Weng, S., Huebner, R.J., Wallingford, J.B., 2022. Convergent extension requires adhesion-dependent biomechanical integration of cell crawling and junction contraction. Cell Reports 39, 110666. 10.1016/j.celrep.2022.110666

Wilson, P.A., Oster, G., Keller, R., 1989. Cell rearrangement and segmentation in Xenopus: direct observation of cultured explants. Development 105, 155–166. 10.1242/dev.105.1.155

Xiong, F., Ma, W., Bénazéraf, B., Mahadevan, L., Pourquié, O., 2020. Mechanical Coupling Coordinates the Co-elongation of Axial and Paraxial Tissues in Avian Embryos. Developmental Cell 55, 354–366.e5. 10.1016/j.devcel.2020.08.007

