## Supplemental figures for "Epithelial-mesenchymal boundary guides cell shapes and axis elongation in embryonic explants"

### Supplementary Figures

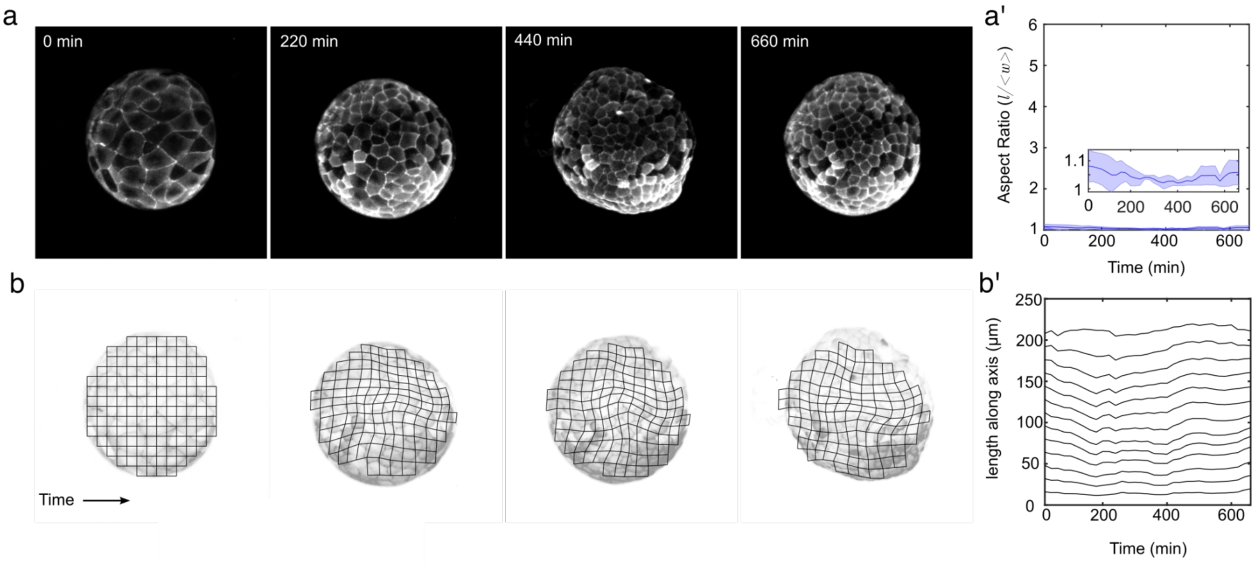


**S1 - Explants without Activin do not elongate**

A) Snapshots of explants with no Activin A') Aspect ratio of explants over time. Scale bar = 100 µm, N=3 B) Tissue deformation over time shows little change. B') Kymograph of horizontal central axis.

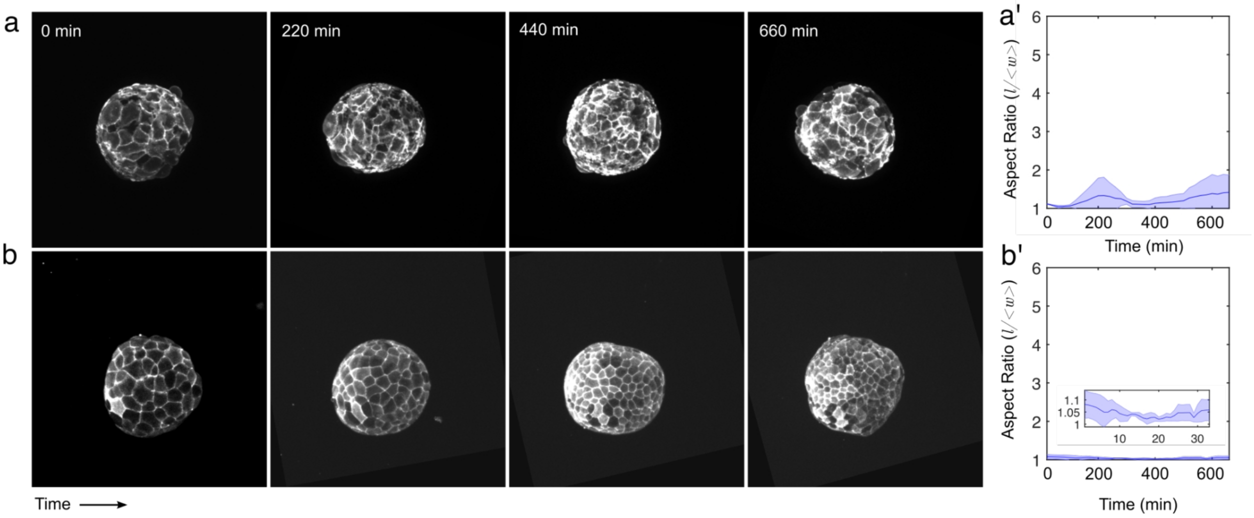


**S2 - Isolated Tissue Experiments**

A) Snapshots of isolated mesenchyme (Mes-) with no Activin. Scale bar = 100µm.  A') Aspect ratio of A, N=3 B) Snapshots of isolated epithelium (Epi-) with no Activin. Scale bar = 100µm. B') Aspect ratio of Epi- experiments, N=3.

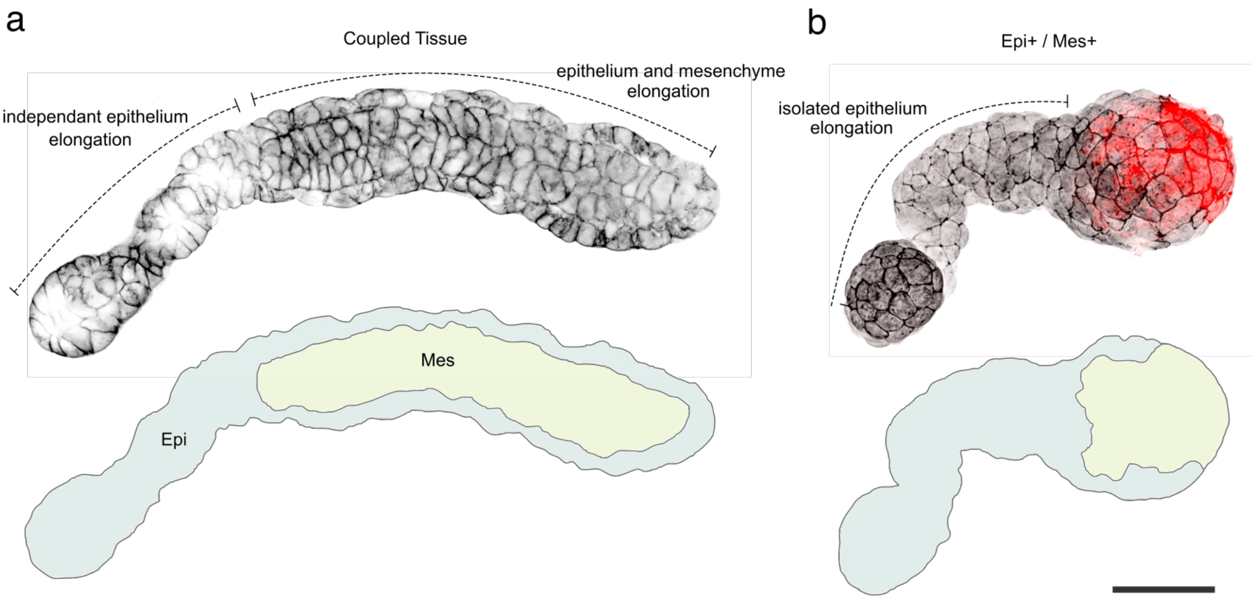


**S3 - The epithelium elongates independently from mesenchymal cells in both intact and when tissues are recombined.**

A) Coupled tissue elongated explant showing internal mesenchymal cells and elongated epithelium without mesenchymal cells. B) Example where Epi+Mes+ recombination experiment shows only the epithelium has extended and the mesenchymal tissue remains spherical.

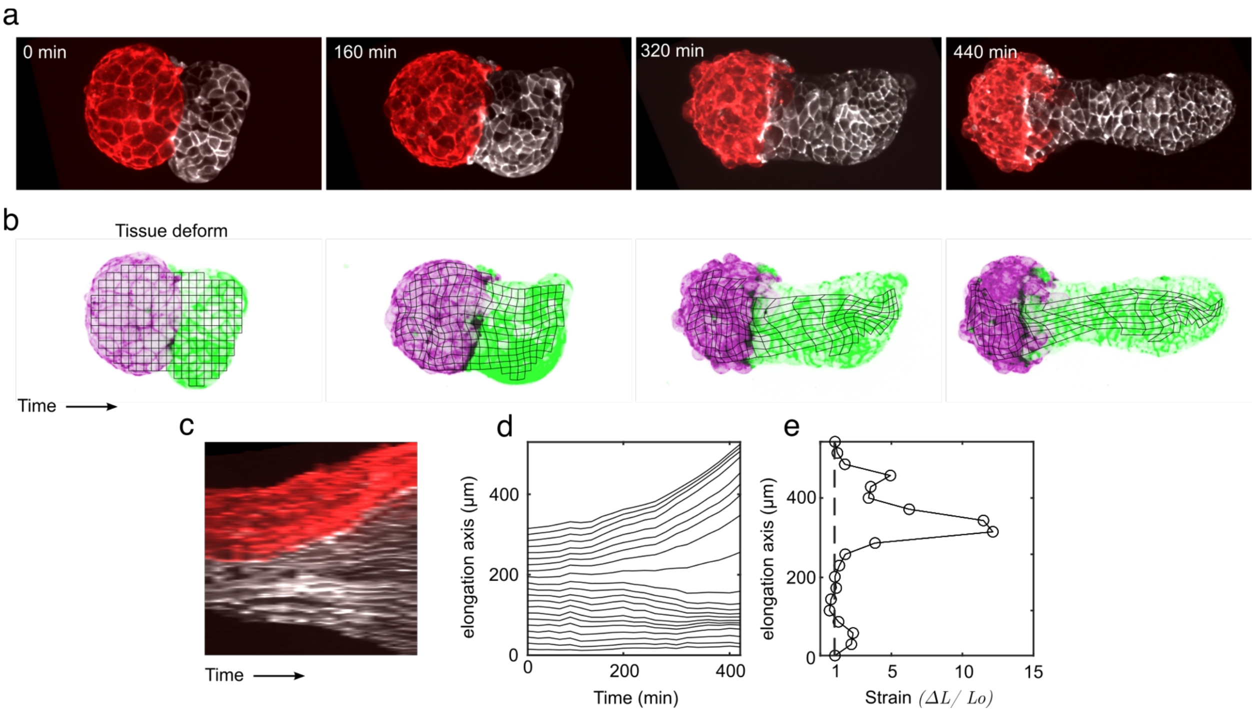


**S4 - Epi+, Mes+ tissue deformation kymograph showing clear tissue elongation at the boundary.**

A)Snapshots of Epi+,Mes+ recombination experiments at different timepoints. B) Tissue deformation at different timepoints, purple is the non-elongating mes and green is the epithelium elongating. C) Kymograph of the explant along the elongating axis D) Tissue deformation kymograph E) Strain of the final timepoint showing the localisation of tissue extension.

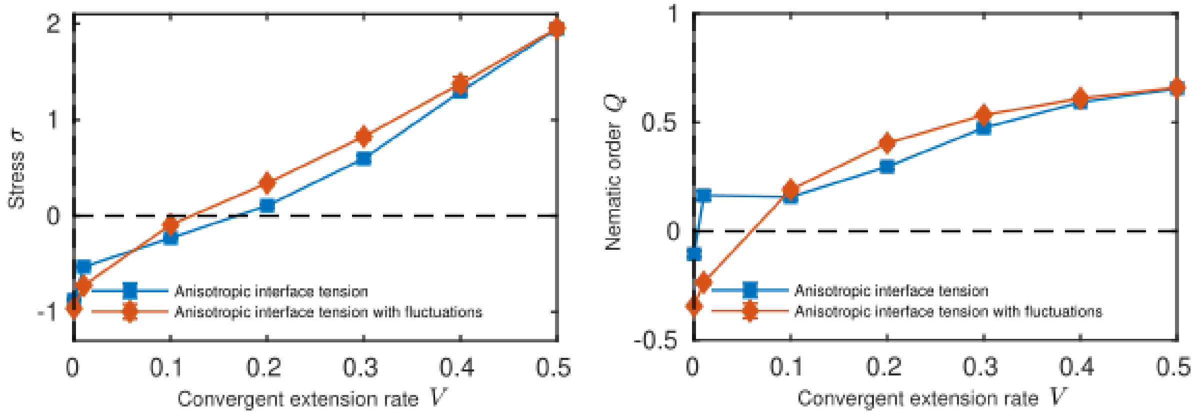


**S5 - Effect of adding fluctuations for active anisotropic interface tension simulations.**
When adding fluctuations to our simulation with active, anisotropic interface tensions we also observe perpendicular alignment of the cell shapes to the axis of tissue elongation. See Methods for parameter values.

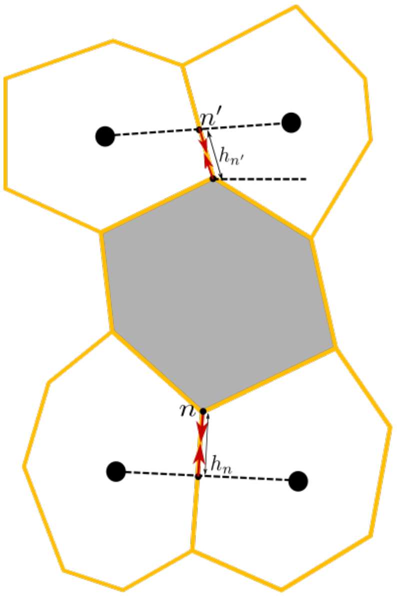


**S6 - Crawl forces are implemented by including active tensions.**
To simulate crawl forces, for each cell $c$ (gray), two active tensions are included in the simulations, one for the vertex with minimal $y$ position and one for the vertex with maximal $y$ position. For any given vertex, the intersection is determined between the interface belonging to the vertex that does not abut cell $c$ and the line connecting the barycenters of the two neighboring cells of $c$ that abut the vertex. A constant active mechanical tension $F_a$ is imposed on the line

connecting the vertex with that intersection point. This approach automatically ensures linear and angular momentum conservation, which is important because there are no relevant external forces acting on the Xenopus explants.
